# Cooperative regulation of NF-E2 related factor 1 protein stability and transcriptional activation by endoplasmic reticulum-associated degradation system mediator, Selenoprotein S/K

**DOI:** 10.64898/2026.05.16.725617

**Authors:** Goki Yamada, Nozomi Tanaka, Yuka Kamada, Reiko U. Yoshimoto, Marino Kita, Hanae Takami, Yudai Suetsugu, Takeshi Sawada, Mizuho A. Kido, Tsukasa Okiyoneda, Tadayuki Tsujita

## Abstract

NRF1 is a key mediator of the proteasome recovery pathway, yet its regulation by ER-resident factors is not fully elucidated. Here, we demonstrate that selenoproteins SELS and SELK are critical regulators for NRF1 protein dynamics. SELS stabilizes NRF1, while SELK induces its insolubilization. Their deficiency leads to a hyper-accumulation and increased nuclear localization of NRF1 under proteasome inhibition condition. This results in an augmented transcriptional response of proteasome subunits. These results indicate that SELS and SELK cooperatively gate NRF1 activity by controlling its retrotranslocation and solubility, highlighting a novel layer of selenoprotein-mediated quality control in the proteostasis network.

## 1. Introduction

The maintenance of cellular protein homeostasis is essential for cell survival and relies heavily on the proper functioning of the ubiquitin-proteasome system. When the proteolytic capacity of the proteasome is impaired by physiological stress or pharmacological inhibitors, cells invoke a compensatory mechanism known as the “bounce-back response” (1). This adaptive process upregulates the de novo synthesis of proteasome subunits and is primarily orchestrated by the NRF1 (NF-E2-related factor 1, also known as NFE2L1), an ER-resident transcription factor of the Cap ‘n’ Collar (CNC) basic region leucine zipper (2).

NRF1 is subjected to a unique and highly complex post-translational regulation. It is initially synthesized as an endoplasmic reticulum (ER) transmembrane protein, with the bulk of its polypeptide residing in the ER lumen. Under homeostatic conditions, NRF1 is rapidly targeted for ER-associated degradation (ERAD); it is ubiquitinated, retrotranslocated to the cytosol by the Valosin-containing protein (VCP), a member of the AAA+ ATPase superfamily, and promptly degraded by the proteasome (3–6). However, upon proteasome inhibition, the retrotranslocated NRF1 escapes degradation. It then undergoes deglycosylation by N-glycanase 1 (NGLY1) (7,8) and subsequent cut off from the ER membrane by the aspartyl protease DNA-damage inducible 1 homolog 2 (DDI2) (9–11). The processed active fragment translocates into the nucleus to transactivate proteasome subunit genes via antioxidant response elements (AREs) (12,13). Thus, for NRF1, the ERAD machinery represents not only a constitutive degradation pathway, but rather an indispensable activation mechanism that extracts the protein into the cytosol to facilitate its nuclear translocation when proteasome activity declines. The indispensable physiological roles of NRF1 have been unequivocally demonstrated by in vivo studies using genetically engineered mouse models. Global disruption of the *Nrf1* gene results in embryonic lethality (14). Furthermore, tissue-specific ablation of *Nrf1* leads to severe localized pathologies; for example, its deletion in the liver causes non-alcoholic steatohepatitis and hepatic neoplasia, while its deficiency in the central nervous system provokes progressive neurodegeneration accompanied by the accumulation of ubiquitinated proteins due to diminished proteasome capacity (15–19).

Elucidating the precise mechanisms underlying NRF1 activation and its subsequent nuclear translocation is of paramount clinical and biological importance, as the NRF1-dependent proteasome regulation represents a promising therapeutic target for multiple human pathologies, including cancer and neurodegenerative diseases. In cancer cells, NRF1 confers resistance to proteasome inhibitors by robustly inducing the bounce-back response; therefore, delineating and therapeutically blocking the pathway that allows NRF1 to escape the ER and enter the nucleus could overcome this drug resistance and potentiate the cytotoxicity of proteasome inhibitors (7,20,21). Conversely, neurodegenerative disorders such as Alzheimer’s, Parkinson’s, and Huntington’s diseases are often characterized by the toxic accumulation of ubiquitinated, aggregate-prone proteins due to diminished proteasome capacity. In these contexts, identifying mechanisms to actively promote the nuclear translocation of NRF1 could offer a viable strategy to enhance de novo proteasome synthesis and accelerate the clearance of toxic aggregates. Despite this immense therapeutic potential, the precise machinery and regulatory factors controlling the dynamic mobilization of NRF1 from the ER to the nucleus remain incompletely understood.

Two ER-resident selenoproteins, SELENOS (also known as SELS or VIMP) and SELENOK (SELK), have been identified as key components of the ERAD multiprotein complex (22,23). SELS functions as a crucial linker by interacting with both DERLIN-1 and VCP, facilitating the extraction of misfolded proteins from the ER to the cytosol. Similarly, SELK binds to the ERAD complex and is intimately involved in maintaining ER homeostasis. Despite possessing no sequence similarity, SELS and SELK share a similar topology, featuring a single transmembrane domain and a cytosolic tail containing a highly reactive selenocysteine (Sec) residue. Recent studies have indicated that SELS and SELK functionally cooperate to form a functional ERAD complex during ER stress. While these selenoproteins have traditionally been considered to simply mediate the clearance of misfolded or denatured proteins, we hypothesized that they might possess an active regulatory mechanism to dynamically control the fate of functional ER membrane proteins. Specifically, we reasoned that SELS and SELK do not simply dispose of aberrant proteins through ERAD, but actively regulate the dynamics of NRF1, such as by suppressing its degradation.

In the present study, we aimed to delineate the roles of SELS and SELK in the regulation of NRF1 stability and its transcriptional activity. We identified that SELS and SELK exert contrasting effects on NRF1: whereas SELS delays the degradation of NRF1 and retains it in the ER, SELK predominantly induces its insolubilization. Furthermore, we demonstrate that the simultaneous loss of SELS and SELK under proteasome impairment leads to the hyper-accumulation of nuclear NRF1, resulting in the excessive transcriptional activation of its target genes. These findings reveal that SELS and SELK are not just redundant ERAD components but rather act cooperatively to fine-tune the basal stability and the stress-induced bounce-back response of endogenous NRF1.

## 2. Materials and methods

### 2.1 Cell line culture condition

Human embryonic kidney (HEK) 293 cell line was obtained from Riken cell bank (Ibaraki, Japan), and cultured in DMEM High glucose (Fujifilm Wako, Osaka, Japan) containing 5% fetal bovine serum (Capricorn scientific, Ebsdorfergrund, Germany), 100 unit/mL penicillin, and streptomycin (Nacalai Tesque, Kyoto, Japan) under humidified air containing 5% CO_2_ at 37°C. Mouse embryonic fibroblast (MEF) was obtained in our laboratory with standard technique (24), and cultured in DMEM High glucose (Fujifilm Wako) containing 5% Newborn Calf Serum (Sigma-Aldrich, St. Louis, MO, USA), 100 unit/mL penicillin, and streptomycin (Nacalai Tesque) under humidified air containing 5% CO_2_ at 37°C. Human embryonic kidney GripTite™ 293MSR cell line (Thermo Fisher Scientific) was cultured in DMEM High Glucose (Fujifilm Wako) supplemented with 10% FBS (Biosera), 0.5 mg/ml G418 (Fujifilm Wako), 100 U/ml penicillin and 100 μg/ml streptomycin (Fujifilm Wako).

### 2.2 Vector construction

The mouse *SelS* and *SelK* cDNA sequences were obtained from multiorgan cDNA library. Both of the cDNAs were introduced into pEF-BOS (25) with FLAG or HA-tag at the N-terminus region. Those vectors were named pEF-BOS-FLAG-SELS (pFLAG-SELS) and pEF-BOS-HA-SELK (pHA-SELK). To monitor the degradation of the NRF1 protein, the HiBiT-tag sequence was inserted into the N-terminus of the pcDNA3.1-NRF1-V5 construct (pNRF1-V5) (26). The resultant vector was named pcDNA3.1-HiBiT-NRF1 (pHiBiT-NRF1). All subcloned sequences were verified by ProDye Terminator sequencing system and Spectrum compact system (Promega, Madison, WI, USA).

### 2.3 Mammalian cell transfection

2.0 × 10^5^ HEK293 cells were plated into 12 well plate and incubated for 16 h. Thereafter, pNRF1-V5, pFLAG-SELS, or pHA-SELK (0.5 µg each) were introduced using Avalanche-Everyday Transfection Reagent (EZ Biosystems, Baltimore, MD, USA). pEF-BOS (empty vector) was used as a mock control and experiments were performed two days after transfection. Transient transfection in 293MSR cells was achieved using polyethylenimine (PEI) max (Polysciences, Warrington, PA, USA). siRNA transfections (50 nM each) were performed using Lipofectamine RNAiMax transfection reagent (Thermo Fisher Scientific). The target sequences of siRNA are listed in Table S1. Steaslth RNAi siRNA Negative Control, Med GC (Thermo Fisher Scientific) was utilized as a negative control non-targeting siRNA (siNT).

### 2.4 Immunoblot analysis

Harvested cells were solubilized with TBS-N buffer (10 mM Tris-HCl, pH 7.5, 150 mM NaCl, 0.5% NP-40) containing a protease inhibitor cocktail (Fujifilm Wako) on ice, and the cell lysates were cleared by centrifugation at 20,000 × g for 10 min at 4°C. The supernatants were mixed with SDS sample buffer (0.25 M Tris-HCl, pH 6.8, 100 mM DTT, 8% [w/v] SDS, 20% [w/v] sucrose, 0.008% [w/v] bromophenol blue) and boiled at 60°C for 10 min. The samples were subjected to SDS-PAGE electrophoresis and transferred to polyvinylidene fluoride membranes (Fujifilm Wako). The blots were incubated with primary antibodies against DDDDK-tag mAb (Medical & biological laboratories, Tokyo, Japan), HA-tag (Merck, Darmstadt, Germany), V5 (Thermo Fisher Scientific), β-Actin (Fujifilm Wako), TCF11/NRF1 (Cell Signaling Technology, Danvers, MA, USA) or α-Tubulin (Fujifilm Wako) and subsequently treated with immunogen-matched horseradish peroxidase-conjugated secondary antibodies. The proteins were visualized with ImmunoStar LD (Fujifilm Wako). The signals were captured using Lumino Graph I (ATTO, Tokyo, Japan).

### 2.5 Immunofluorescence

HEK293 cells and MEF cells were fixed with 4% formaldehyde for 10 min, washed three times with 1 × PBS, and permeabilized with 0.1% Triton X-100 in 1 × PBS for 10 min. The cells were washed three times with 1 × PBS and treated with the primary antibodies: anti-DDDDK-tag mAb, anti-V5, anti-Calnexin (Cell Signaling Technology), anti-HA, anti-TCF11/NRF1 (Cell Signaling Technology) for 1 h at room temperature. Cells were washed with T-TBS (20 mM Tris-HCl, pH 8.0, 150 mM NaCl, 0.1% Tween 20) three times and were incubated with the secondary antibodies: anti-mouse IgG DyLight 488 (Bio-Rad, Hercules, CA, USA), Anti-rabbit IgG Alexa Fluor 594 conjugate (Cell Signaling Technology), anti-rabbit IgG DyLight488 (Bio-Rad) for 1 h at room temperature. After cells were washed three times with T-TBS, the nuclei were stained with 4’, 6-diamidino-2-phenylindole dihydrochloride (DAPI, Bio-Rad). The samples were washed with 1 × PBS, Fluoro-KEEPER Antifade regent (Nacalai Tesque) was placed on the glass slides. Fluorescence images were captured by LSM 880 confocal microscope equipped with Airlyscan (ZEISS, Oberkochen, Germany)

### 2.6 HiBiT degradation assay

For SELS/K overexpression, subconfluent 293MSR cells in six-well plates were transfected with pFLAG-SELS, pHA-SELK, or both pFLAG-SELS and pHA-SELK, along with pHiBiT-NRF1 and cytosolic LgBiT (pBiT1.1-N [TK/LgBiT], Promega). The next day, cells were trypsinized and seeded in 96-well plates (2.0 × 10^4^ cells/well) and cultured for 18–24 h. Cells were washed with 100 μL of the CO_2_-independent medium (Thermo Fisher Scientific). Then, 50 μL of 0.1 × Nano-Glo Endurazine (Promega) in the CO_2_-independent medium was added to each well. The cells were incubated for 2.5 h at 37°C and 5% CO_2_.

For SELS/K knockdown, subconfluent 293MSR cells in six-well plates were transfected with siRNA (50 nM each). After one day of culture, the transfectant were trypsinized, reseeded into six-well plates, and cultured for another day. Then, the cells were transfected with pHiBiT-NRF1 and cytosolic LgBiT. The following day, the cells were trypsinized again and seeded in Nunc MicroWell 96-Well Nunclon Delta-Treated Flat-Bottom Microplates (Thermo Fisher Scientific) and cultured for 18–24 h. After washing the cells with 100 μL of CO_2_-independent medium, 50 μl of 0.1 × Nano-Glo Endurazine in CO_2_-independent medium was added to each well, and the cells were incubated for 2.5 h at 37°C in 5% CO_2_.

To measure the degradation kinetics, 10 μL of 600 µg/mL Cycloheximide (CHX, Fujifilm Wako) was added to each well, resulting in a final concentration of 100 µg/mL. Luminescence was continuously measured at 5-min intervals for 3 h using a Luminoskan plate reader (Thermo Fisher Scientific). The luminescence signal of CHX-treated cells (LumCHX) was normalized to the signal of untreated cells (Lumuntreat) to calculate the remaining ERAD substrates during the CHX chase as Eq. 1. The ERAD rate was calculated by fitting the luminescence decay with a one-phase exponential decay function using GraphPad Prism 10 (GraphPad, San Diego, CA, USA).

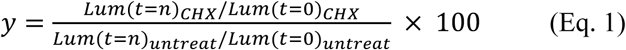

### 2.7 Generation of SELS, SELK single knockout and, SELS/K double knockout cells

CRISPR-Cas9 mediated gene knockout of SELS or SELK (SKO and KKO, respectively) was performed using the pSpCas9(BB)-2A-GFP (PX458) (#48138; Addgene, Watertown MA, USA) containing SELS gRNA (5’–CAC CGT GCC GCT TAA CAA CAA CAT C–3’), SELK gRNA (5’–CAC CGT AGG TCA GGT GTT GGA CAG C–3’). Pairs of oligonucleotides with BbsI (New England Biolabs, Ipswich, MA, USA) overhangs were annealed and ligated into the BbsI-digested vector (PX458-SELS, PX458-SELK). All constructs were verified by ProDye Terminator sequencing system and Spectrum compact system (Promega). For CRISPR/Cas9-mediated knockouts, MEF cells were transfected with PX458-SELS or PX458-SELK using Neon Transfection System (Thermo Fisher Scientific). The transfectants were harvested and suspended into 1 × PBS containing 0.5% BSA and 2.0 mM EDTA. The viable cell harbored GFP fluorescence were sorted individually into 96 well plate with Cell Sorter MA900 (Sony, Tokyo, Japan). SELS/K double knockout (DKO) MEF cells were generated introducing PX458-SELK to SELS knockout MEF cells. Designated SELS/K double knockout MEF cells were selected with same strategy for SELS or K single knockout.

### 2.8 Immunoprecipitation

1.0 × 10^6^ WT-, SKO-, KKO-, DKO-MEF cells were plated into 10 cm dish and incubated for 16 h. The cells were solubilized with 1.0 mL IP buffer (50 mM Tris-HCl, pH 7.5, 100 mM NaCl, 1.0% Triton X-100, 1.0 mM EDTA) containing a protease inhibitor cocktail (Fujifilm Wako), incubated on ice for 30 min, and then centrifuged at 18,000 × g for 20 min at 4°C. The resulting supernatants were incubated with 0.2 µg of anti-TCF11/NRF1 antibody overnight at 4°C with rotation. Subsequently, Ab-Capcher MAG2 beads (ProteNova, Kagawa, Japan) were added to the mixture, and the incubation was continued overnight at 4°C with rotation. The beads were then washed twice with 100 µL of 1 × PBS, and the bound proteins were eluted twice using 15 µL of 0.1 M Glycine-HCl (pH 2.7), yielding a total volume of 30 µL. The eluates were neutralized by adding 1.0 µL of 1 M Tris, mixed with 10 µL of 4 × sample buffer, and incubated at 60°C for 10 min. The resulting samples were then subjected to immunoblot analysis.

### 2.9 RNA isolation and real-time qPCR

4.3 × 10^5^ WT- and DKO-MEF cells were plated in 6 well plate and incubated for 16 h. Thereafter, the MEF cells were treated with 1.0 µM MG132 for 24 h. Total RNA was prepared from the MEF cells using Sepasol-RNA I Super G (Nacalai Tesque) according to the manufacturer’s instructions. A 1-µg aliquot of the total RNA was reverse transcribed using Access RT-PCR System (Promega). The cDNA was used as a template for qRT-PCR using KAPA SYBR FAST qPCR Master Mix system (Kapa Biosystems, Wilmington, MA, USA). The qPCR primers for *Nrf1*, *Psmb7* and *18S* are listed in Table S2.

## 3. Results

### 3.1 SELS overexpression increases NRF1 protein levels by retaining it in the ER

To investigate the regulatory role of SELS for NRF1 protein stability, we first performed co-overexpression experiments by transfecting pFLAG-SELS and pNRF1-V5 into HEK293 cells. NRF1-V5 protein levels were significantly increased when we co-overexpressed the FLAG-SELS protein. The accumulated NRF1-V5 protein, which appeared as 140K molecular weight signal, exhibited a downward shift when treated with PNGase F. This shift confirms that the accumulated protein represents an N-glycosylated form of NRF1(Fig. 1A) (27). Subsequent immunofluorescence analysis revealed that FLAG-SELS localized to the endoplasmic reticulum (ER). Notably, we observed that the ER in these FLAG-SELS-overexpressing cells underwent morphological reorganization, forming filamentous ER-microtubule (MT) bundles (Fig. 1B). This observation is consistent with a previous report demonstrating that SELS physically links the ER to MTs, and its overexpression robustly induces ER-MT bundling (28,29). Furthermore, we examined the localization of NRF1-V5 in these cells and found that the accumulated NRF1-V5 predominantly colocalized with the ER marker protein CALNEXIN (Fig. 1C). These findings suggest that SELS suppresses the retrotranslocation or degradation of NRF1, thereby retaining it within the ER membrane.

**Figure 1.**
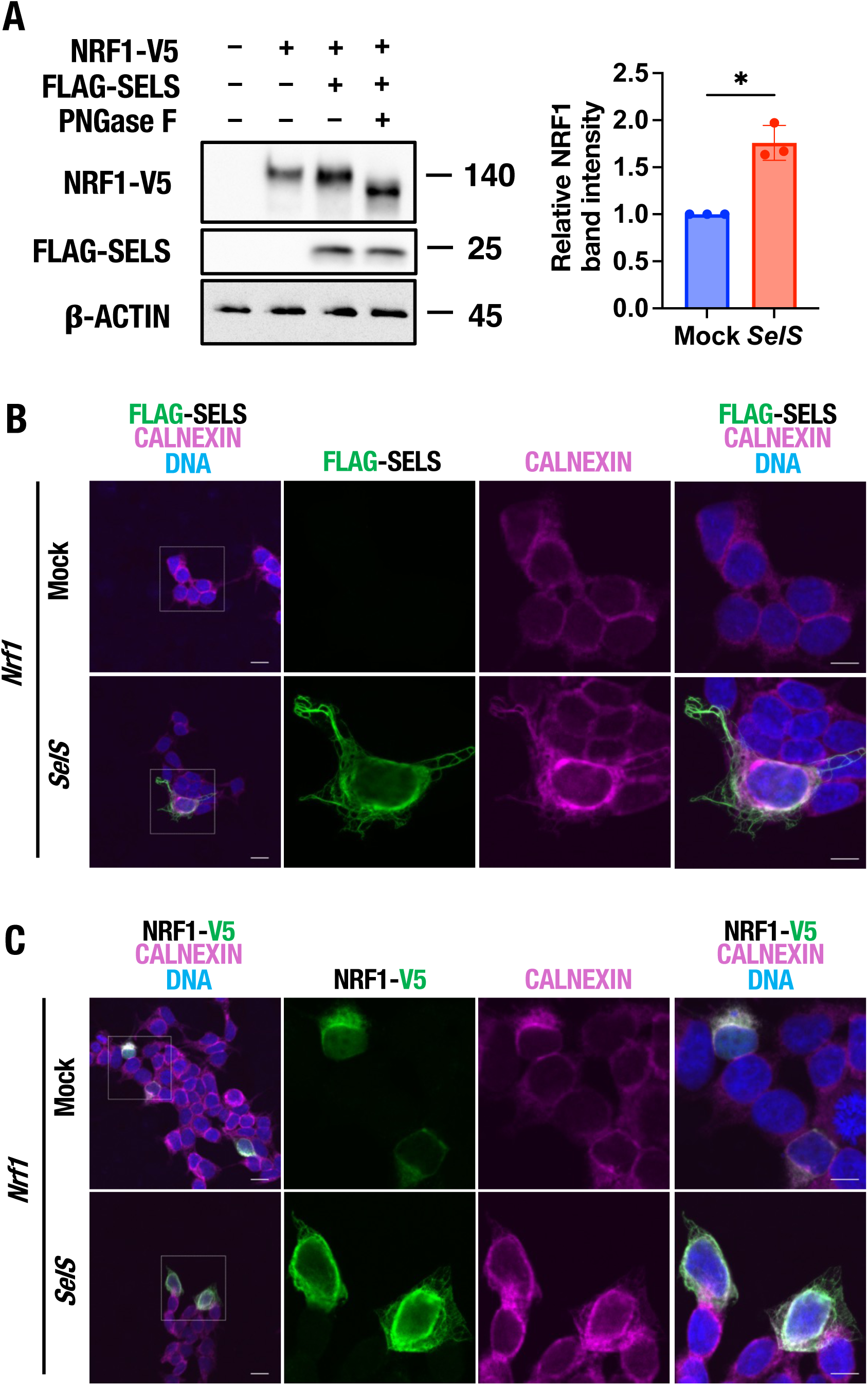
SELS and NRF1 co-overexpression induces an accumulation of N-glycosylated NRF1 on ER membrane. **(A) Accumulation of N-glycosylated NRF1 upon SELS overexpression.** HEK293 cells were transfected with an empty vector (Mock), pNRF1-V5 alone, or co-transfected with pNRF1-V5 and pFLAG-SELS. Cell lysates from the co-transfected cells were further treated with or without PNGase F. The samples were analyzed by immunoblotting using anti-V5 and anti-FLAG antibodies (left). The graph shows the quantification of NRF1-V5 protein levels (right). The NRF1-V5 protein levels are normalized with ß-ACTIN. The accumulated NRF1-V5 protein, appearing at approximately 140K in FLAG-SELS-overexpressing cells. The downward shift upon PNGase F treatment, confirming its N-glycosylated status. Band intensities were quantified using ImageJ Fiji software. Data are presented as mean ± SD of three independent experiments (n = 3). Statistical significance was determined using a paired Student’s *t*-test (\**P* < 0.05). **(B) Subcellular localization of FLAG-SELS.** Immunofluorescence analysis of HEK293 cells expressing NRF1-V5 alone (top row) or co-expressing NRF1-V5 and FLAG-SELS (bottom row). Cells were immunostained for FLAG (SELS), CALNEXIN (ER), and nucleic DNA. The leftmost panels display merged images at 400× magnification. From left to right, the remaining panels, show 1,260× magnified images of the FLAG signal, calnexin signal, and the merged signals, respectively. Scale bars represent 20 µm in 400× images, and 10 µm in 1,260× images. **(C) ER retention of NRF1 in SELS-overexpressing cells.** Immunofluorescence analysis of HEK293 cells under the same transfection conditions as in (B). Cells were immunostained for V5 (NRF1), CALNEXIN, and nucleic DNA. The panel layout is identical to (B), showing 400× merged images on the far left, followed by 1260× magnified images of the V5, CALNEXIN, and merged signals. Scale bars represent 20 µm in 400× images and 10 µm in 1,260× images.

### 3.2 SELK overexpression decreases NRF1 protein levels

We next examined the effect of SELK, another ER-resident selenoprotein closely related to SELS, on NRF1. In striking contrast to SELS, co-overexpression of HA-SELK resulted in a marked decrease in NRF1-V5 protein levels in HEK293 cells as detected by immunoblotting (Fig. 2A). Subsequent immunofluorescence analysis confirmed that HA-SELK localized to the ER, overlapping with the ER marker CALNEXIN (Fig. 2B). Notably, unlike SELS, HA-SELK overexpression did not induce drastic morphological changes such as ER-MT bundling, maintaining a typical ER network structure. Furthermore, despite the apparent substantial loss of NRF1-V5 in the immunoblot analysis, our immunofluorescence imaging clearly revealed that NRF1-V5 was still retained within the ER in SELK-overexpressing cells (Fig. 2C). This intriguing discrepancy between the biochemical and morphological observations implies that SELK does not simply promote the complete degradation and clearance of NRF1 from the cell. Instead, it suggests that SELS and SELK exert fundamentally distinct modes of regulation on the physical state of NRF1.

**Figure 2.**
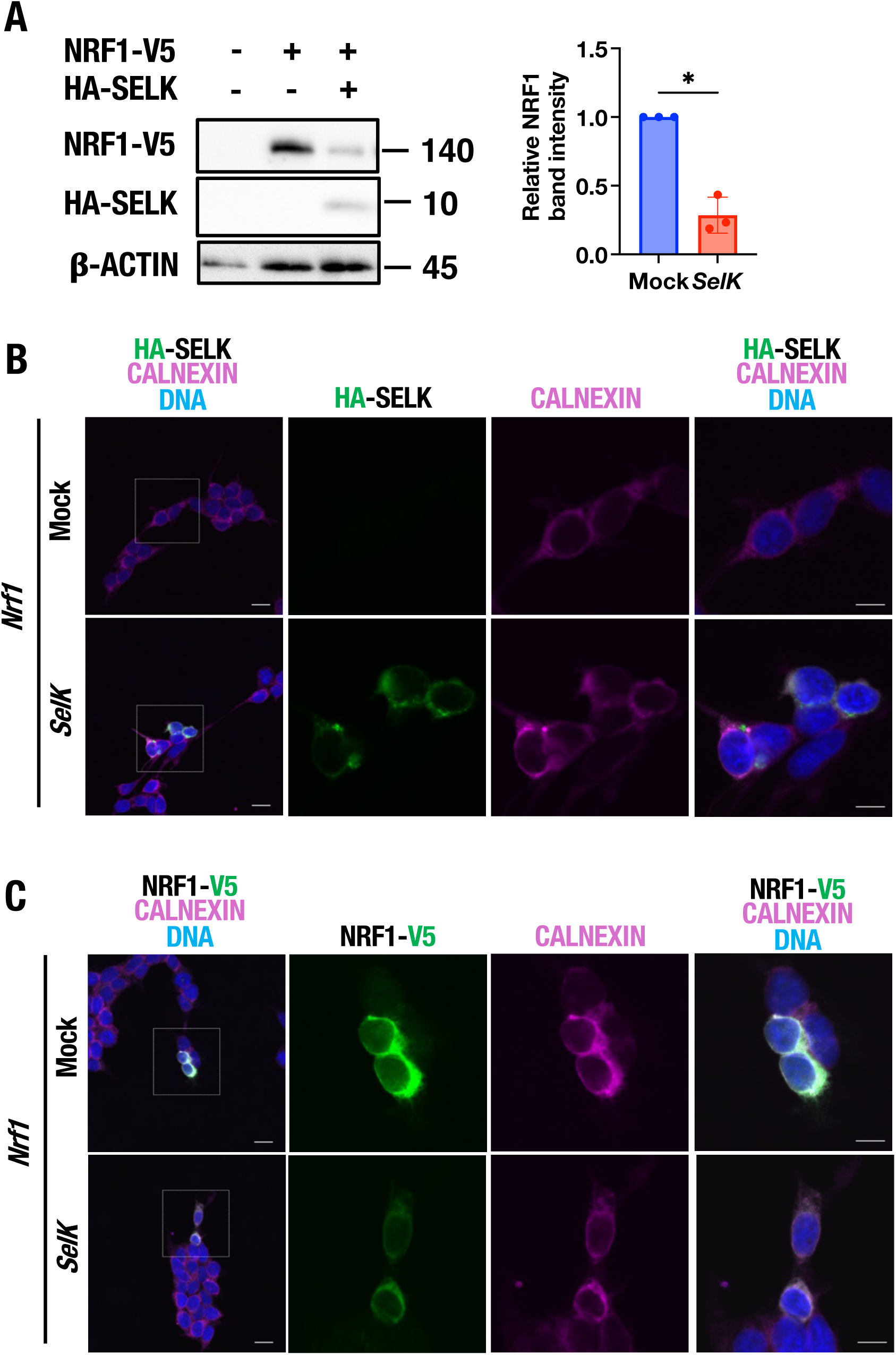
SELK and NRF1 co-overexpression induces a decrease in NRF1 on ER membrane. **(A) Reduction of N-glycosylated NRF1 upon SELK overexpression.** HEK293 cells were transfected with an empty vector (Mock), pNRF1-V5 alone, or co-transfected with pNRF1-V5 and pHA-SELK. The samples were analyzed by immunoblotting using anti-V5 and anti-HA antibodies (left). The graph shows the quantification of NRF1-V5 protein levels (right) The NRF1-V5 protein levels are normalized with ß-ACTIN. Band intensities were quantified using ImageJ Fiji software. Data are presented as mean ± SD of three independent experiments (n = 3). Statistical significance was determined using a paired Student’s *t*-test (\**P* < 0.05). **(B) Subcellular localization of HA-SELK.** Immunofluorescence analysis of HEK293 cells expressing NRF1-V5 alone (top row) or co-expressing NRF1-V5 and HA-SELK (bottom row). Cells were immunostained for HA (SELK), CALNEXIN (ER), and nucleic DNA. The leftmost panels display merged images at 400× magnification. From left to right, the remaining panels show 1,260× magnified images of the FLAG signal, calnexin signal, and the merged signals, respectively. Scale bars represent 20 µm in 400× images and 10 µm in 1,260× images. **(C) Subcellular localization of NRF1 in SELK-overexpressing cells.** Immunofluorescence analysis of HEK293 cells under the same transfection conditions as in (B). Cells were immunostained for V5 (NRF1), CALNEXIN, and nucleic DNA. The panel layout is identical to (B), showing 400× merged images on the far left, followed by 1,260× magnified images of the V5, CALNEXIN, and merged signals. Scale bars represent 20 µm in 400× images and 10 µm in 1,260× images.

### 3.3 SELS delays NRF1 degradation, whereas SELK induces its insolubilization

To further elucidate the dynamics of NRF1 regulation by SELS and SELK, we quantitatively measured the degradation rate of NRF1 using a HiBiT degradation assay (Fig. 3A). In this assay, HiBiT-NRF1 was co-expressed with a cytosol-localized LgBiT in 293 MSR cells. Following the arrest of de novo protein synthesis with cycloheximide (CHX), the luminescent signal decay was continuously monitored. Consistent with the immunoblot results, SELS overexpression significantly delayed the decay of the luminescent signal, indicating that SELS decreases the degradation rate of NRF1 (Fig. 3B).

**Figure 3.**
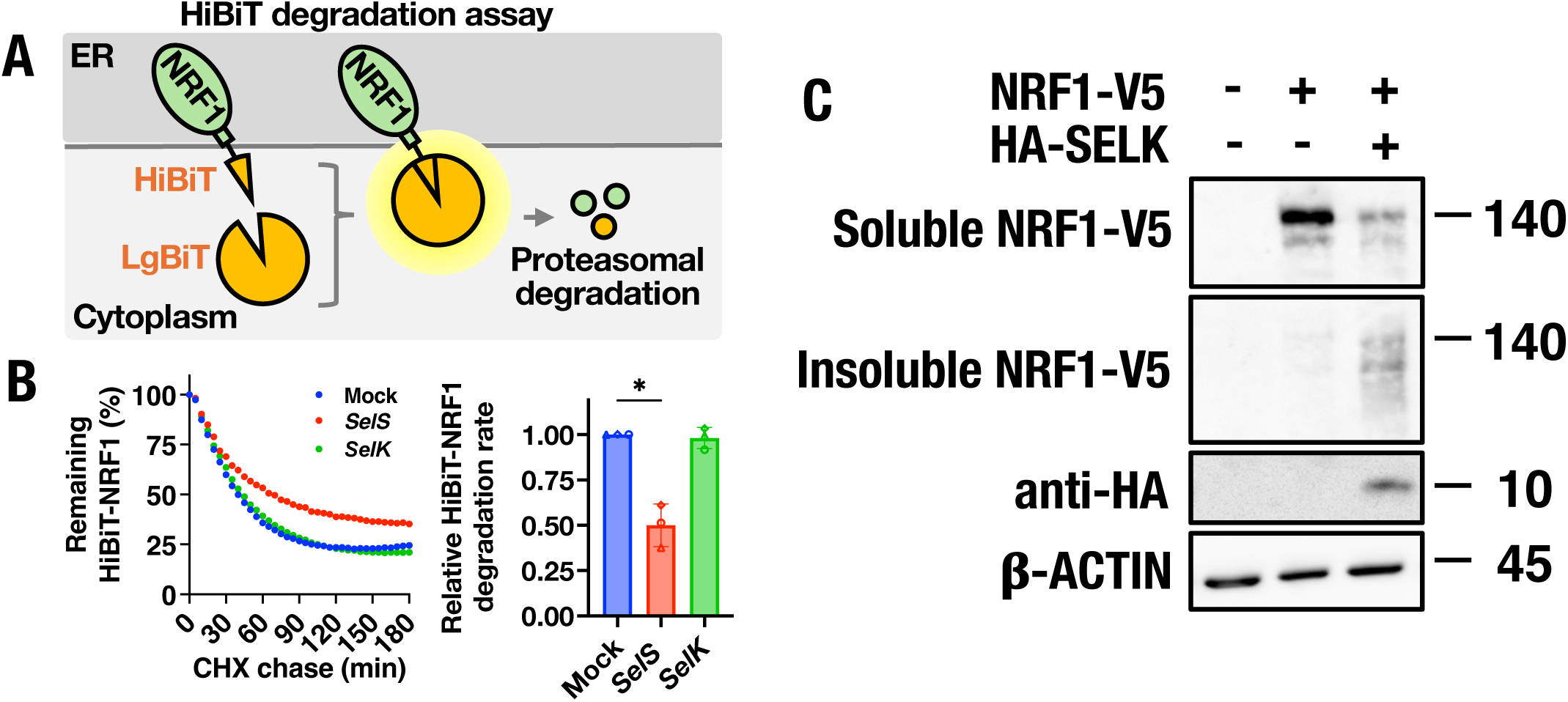
SELS delays NRF1 degradation, whereas SELK induces its insolubilization. **(A) Schematic representation of the HiBiT degradation assay.** HiBiT-tagged NRF1 (HiBiT-NRF1) is co-expressed with a cytosol-localized LgBiT. The binding of HiBiT to LgBiT reconstitutes a functional nano luciferase, enabling luminescence detection in the presence of a substrate. As HiBiT-NRF1 undergoes proteasomal degradation, the luminescent signal proportionally decreases. **(B) SELS overexpression decreases the degradation rate of NRF1.** GripTite 293 MSR cells expressing HiBiT-NRF1 and LgBiT were co-transfected with an empty vector (Mock), pFLAG-SELS, or pHA-SELK. Following the arrest of de novo protein synthesis with cycloheximide (CHX), luminescence decay was continuously monitored for 180 min (left). The graph shows the quantification of the relative HiBiT-NRF1 degradation rate (right). Data are presented as mean ± SD of three independent experiments (n = 3). Statistical significance was determined using a one-way repeated-measures (RM) ANOVA with Dunnett’s multiple comparison test (\**P* < 0.05). **(C) Accumulation of insoluble NRF1 in SELK-overexpressing cells.** HEK293 cells were transfected with pNRF1-V5 alone or co-transfected with pNRF1-V5 and pHA-SELK. Cell lysates were sonicated and subsequently centrifuged at 15,000 × g for 15 min. The resulting supernatants were collected as the soluble fraction, whereas the pellets were resuspended in the 1× SDS sample buffer to obtain the insoluble fraction. These fractions are denoted as Soluble NRF1-V5 and Insoluble NRF1-V5, respectively. The samples were then analyzed by immunoblotting.

A previous study has reported that treatment with high concentrations of proteasome inhibitors causes NRF1 to aggregate and disappear from soluble cell lysates. Furthermore, it has been demonstrated that such aggregation induced by high concentrations of proteasome inhibitors concurrently suppresses the transcriptional activity of NRF1 (30,31). Based on this finding, we attempted to detect insoluble NRF1 by fractionating cell lysates into detergent-soluble and - insoluble fractions. Given the apparent reduction of NRF1 upon SELK overexpression (Fig. 2A), we hypothesized that SELK might induce the insolubilization of NRF1 rather than simply promoting its complete degradation. To test this, we fractionated cell lysates into detergent-soluble and -insoluble fractions. Strikingly, while NRF1-V5 was typically found in the soluble fraction, HA-SELK overexpression caused an accumulation of NRF1-V5 in the insoluble fraction (Fig. 3C). These results indicate that SELS stabilizes NRF1 by delaying its degradation, whereas SELK predominantly induces its insolubilization.

### 3.4 SELS and SELK cooperatively regulate endogenous NRF1 stability and its transcriptional activation

To examine how the reduction of SELS and SELK protein levels affects NRF1 stability, we performed siRNA-mediated knockdown of SELS and SELK, which was confirmed by Immunoblot analysis (Fig. 4A). HiBiT degradation assays revealed that the depletion of SELK increased the degradation rate of NRF1 (Fig. 4B). Furthermore, simultaneous knockdown of both SELS and SELK (double KD) resulted in a significant NRF1 degradation. These results suggest that endogenous SELS and SELK cooperatively contribute to the maintenance of NRF1 stability. We subsequently evaluated the cooperative role of endogenous SELS and SELK in NRF1 regulation. Using the HiBiT degradation assay, we found that siRNA-mediated knockdown of SELK significantly accelerated NRF1 degradation, and simultaneous knockdown of SELS and SELK (si*SelS/K*) further accelerated this process (Fig. 4A). The knockdown efficiencies of SELS and SELK were confirmed by immunoblotting (Supplemental Fig. 1A).

**Figure 4.**
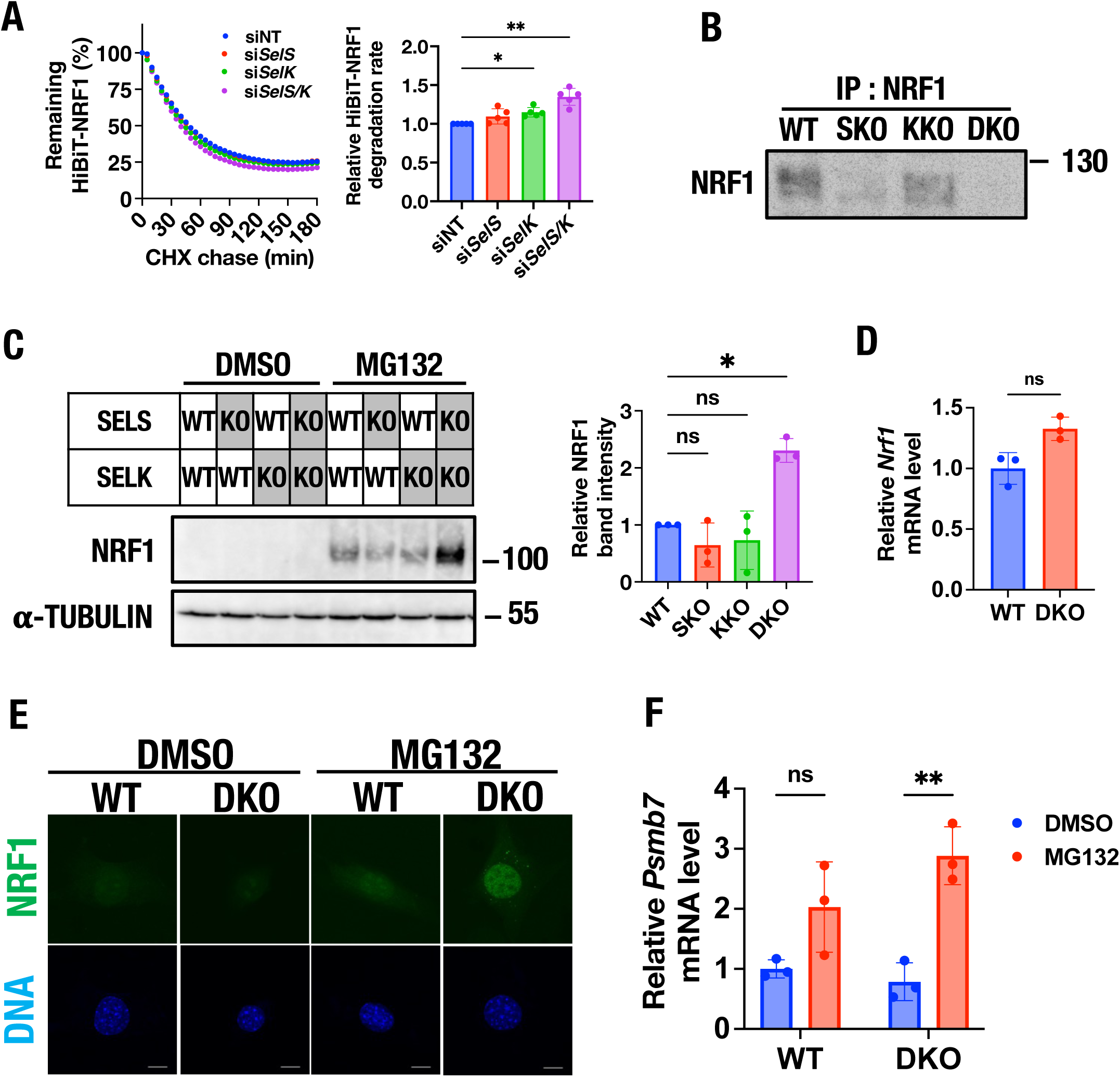
SELS and SELK cooperatively regulate endogenous NRF1 stability and its transcriptional activation. **(A) Accelerated NRF1 degradation upon SELK or simultaneous SELS/SELK knockdown.** GripTite 293 MSR cells expressing HiBiT-NRF1 and LgBiT were transfected with non-targeting control siRNA (siNT), *SelS* siRNA (si*SelS*), *SelK* siRNA (si*SelK*), or both *SelS* and *SelK* siRNAs (si*SelS/K*). Following the arrest of de novo protein synthesis with cycloheximide (CHX), luminescence decay was continuously monitored (left). The graph shows the quantification of the HiBiT-NRF1 degradation rate (right). Data are presented as mean ± SD of five independent experiments (n = 5). Statistical significance was determined using a one-way repeated-measures (RM) ANOVA with Dunnett’s multiple comparison test (\**P* < 0.05, \*\**P* < 0.01). **(B) Decreased basal levels of endogenous NRF1 in SELS/SELK double knockout cells.** Endogenous NRF1 protein levels in wild-type (WT), SELS knockout (SKO), SELK knockout (KKO), and SELS/SELK double knockout (DKO) mouse embryonic fibroblasts (MEFs) were analyzed. Endogenous NRF1 was enriched by immunoprecipitation using an anti-NRF1 antibody prior to immunoblot analysis. **(C) Hyper-accumulation of NRF1 in DKO cells following proteasome inhibition.** WT, SKO, KKO, and DKO MEFs were treated with 0.1% DMSO or 1.0 µM MG132 for 12 h. Cell lysates were analyzed by immunoblotting (left). The graph shows the quantification of endogenous NRF1 protein levels (right). Data are presented as mean ± SD of three independent experiments (n = 3). Statistical significance was determined using a repeated-measures (RM) one-way ANOVA with a Dunnett’s multiple comparison test (\**P*< 0.05). **(D) NRF1 mRNA levels remain unchanged in DKO cells.** The relative mRNA levels of *Nrf1* in WT and DKO MEFs were quantified by qPCR. Data are presented as mean ± SD of technical triplicates from a single experiment. Statistical significance was determined using a paired Student’s *t*-test (no significant difference). **(E) Enhanced nuclear accumulation of NRF1 in DKO cells during proteasome inhibition.** Immunofluorescence analysis of endogenous NRF1 in WT and DKO MEFs are treated with 0.1% DMSO or 0.5 µM MG132 for 24 h. Cells were immunostained for NRF1 (top row) and nucleic DNA (bottom row). All images are shown at 1,260x magnification. Scale bars represent 10 µm. **(F) Hyperactivation of NRF1-target gene transcription in DKO cells.** The relative mRNA levels of the proteasome subunit gene *Psmb7* in WT and DKO MEFs are quantified by qPCR following treatment with 0.1% DMSO or 1.0 µM MG132 for 24 h. Data are presented as mean ± SD of technical triplicates from a single experiment. Statistical significance was determined using a two-way ANOVA with Šídák’s multiple comparisons test (\*\**P* < 0.01).

To confirm these findings, we knocked out SELS and SELK in MEF cells using CRISPR-Cas9 system and generated SELS knockout (SKO), SELK knockout (KKO) and SELS/K double KO (DKO) MEF cells. The successful depletion of SELS and SELK in the single or double knockout (DKO) MEFs, as well as the genomic mutations introduced by the CRISPR-Cas9 system, were verified (Supplemental Fig. 1B and C). Endogenous NRF1, enriched by immunoprecipitation, exhibited decreased basal levels specifically in SELS/SELK double knockout (DKO) MEFs (Fig. 4B). Because the mRNA levels of *Nrf1* remained unchanged in DKO cells (Fig. 4D), this reduction reflects post-transcriptional destabilization.

NRF1 is known to mediate the “bounce-back” response by upregulating proteasome genes upon proteasome inhibition. Interestingly, when cells were treated with the proteasome inhibitor MG132, endogenous NRF1 excessively accumulated in DKO cells compared to WT cells (Fig. 4C). Immunofluorescence analysis further revealed a marked enhancement of nuclear accumulation of NRF1 in MG132–treated DKO cells (Fig. 4E). Consequently, the hyper-accumulation of nuclear NRF1 in DKO cells led to the hyperactivation of its transcriptional target; the mRNA expression of the proteasome subunit gene *Psmb7* was significantly higher in DKO cells than in WT cells following MG132 treatment (Fig. 4F). Collectively, these data suggest that the loss of SELS and SELK may accelerate NRF1 turnover, their absence leads to an exaggerated NRF1-mediated transcriptional response when the proteasome is compromised, likely due to altered protein dynamics or quality control.

## 4. Discussion

In the present study, we demonstrated that the ER-resident selenoproteins SELS and SELK cooperatively regulate the steady-state stability and stress-induced transcriptional activation of NRF1. While previous studies have primarily characterized SELS and SELK as components of the ERAD machinery responsible for clearing misfolded proteins, our findings propose a novel concept: these ERAD factors actively protect and fine-tune functional membrane proteins. By exerting contrasting effects—SELS promoting ER retention and SELK inducing insolubilization upon overexpression—they orchestrate the precise dynamics of NRF1, ensuring an appropriate “bounce-back” response during proteasome impairment.

The contrasting effects of SELS and SELK on NRF1 stability highlight their distinct modes of action. At first glance, SELS overexpression might be expected to accelerate NRF1 degradation by excessively recruiting VCP to the ER, given that SELS is known to promote the degradation of misfolded proteins such as the cystic fibrosis transmembrane conductance regulator (CFTR) ΔF508 mutant (32). However, we observed that SELS overexpression paradoxically led to NRF1 accumulation. We hypothesize that overabundant SELS might competitively sequester a large portion of the cellular VCP pool, thereby depleting the available VCP for other ERAD substrates. Consequently, VCP is restricted from accessing NRF1, preventing its proper retrotranslocation and thereby retaining NRF1 within the ER membrane.

Conversely, SELK overexpression led to a marked decrease in soluble NRF1 levels, coupled with its accumulation in the insoluble fraction. It is well documented that SELK serves as an essential cofactor for the ER-localized palmitoyltransferase DHHC6. During this process, the highly reactive Sec residue of SELK is critical for stabilizing the acyl-DHHC6 intermediate to efficiently catalyze the palmitoylation of target proteins (33,34). Based on these lines of evidence, we hypothesize that excessive SELK might drive aberrant DHHC6-mediated palmitoylation of NRF1 or its interacting partners, directly triggering the physical insolubilization or aggregation of NRF1. Future studies focusing on how the Sec residue of SELK specifically affects NRF1 stability will be necessary to fully elucidate this mechanism.

Furthermore, our data from SELS/SELK double knockout (DKO) cells underscore the physiological necessity of this cooperative regulation. Under basal conditions, DKO cells exhibited accelerated NRF1 degradation, suggesting that SELS and SELK ordinarily suppress the premature degradation of NRF1, maintaining a readily available pool of this transcription factor at the ER. Strikingly, upon proteasome inhibition with MG132, DKO cells showed a hyper-accumulation of nuclear NRF1 and an excessive upregulation of the proteasome subunit gene *Psmb7*. This hyperactivation implies that without the regulatory restraint provided by SELS and SELK, the ER-to-nucleus mobilization of NRF1 becomes uncoupled from normal homeostatic checkpoints. Therefore, for NRF1, the SELS/SELK-mediated ERAD complex serves not merely as a degradation apparatus, but as an active gatekeeper that dictates whether NRF1 should be degraded, retained, or fully retrotranslocated for nuclear signaling.

Interestingly, previous in vivo studies have demonstrated that mice globally lacking either SELS or SELK are viable and exhibit relatively mild phenotypes under basal conditions. For instance, SELS knockout mice develop normally but display subtle impairments in fast-twitch muscle contractile function (35), while SELK knockout mice are healthy but exhibit deficient calcium flux and impaired immune responses in leukocytes (36). These mild and restricted phenotypes of single knockouts contrast starkly with the severe dysregulation of NRF1 we observed in our SELS/SELK double knockout (DKO) cells. This discrepancy strongly suggests that SELS and SELK functionally compensate for each other to maintain crucial ER proteostasis and NRF1 dynamics. Therefore, we hypothesize that the simultaneous loss of both SELS and SELK in vivo would completely abolish this compensatory fail-safe mechanism, precipitating a catastrophic breakdown of the NRF1-dependent stress response. Given that global NRF1 deletion is embryonic lethal, it is highly probable that a SELS/SELK double knockout mouse would suffer from severe systemic pathologies or profound developmental defects. Generating and characterizing such DKO animal models in future studies will be an essential next step to fully elucidate the physiological imperative of this cooperative selenoprotein network in living organisms.

Furthermore, the functional interplay between NRF1, SELS, and SELK provides a tantalizing glimpse into an intricate feedback loop maintaining cellular selenium and redox homeostasis. Under basal conditions, NRF1 is known to act as a transcriptional repressor for specific genes, including the cystine/glutamate antiporter *xCT* and the SELENOP receptor *ApoER2* (*LRP8*), thereby restricting excessive antioxidant uptake and selenium influx (15,37,38). We hypothesize that this NRF1-mediated repression is tightly monitored by the Sec-dependent functions of SELS and SELK. When intracellular selenium becomes limiting, the efficiency of Sec insertion at the UGA codon drops, leading to the generation of truncated, Sec-lacking SELS and SELK. Recent studies have revealed that these non-functional, incomplete selenoproteins are rapidly recognized and degraded by specific ubiquitin ligases such as KLHDC1 and CRL2 (28,39). The consequent depletion of functional SELS and SELK at the ER membrane would fundamentally alter NRF1 dynamics. The loss of NRF1’s repressive function would lead to the robust derepression and up-regulation of *xCT* and *LRP8*. This adaptive response would drive a massive influx of cystine and SELENOP, rapidly replenishing the cellular redox buffering capacity and selenium pool to support the de novo synthesis of functional selenoproteins. Thus, SELS and SELK may function as selenium-dependent modulators of NRF1 behavior, connecting cellular selenium availability to a vast homeostatic network that links ER proteostasis, redox balance, and trace element metabolism.

The elucidation of this SELS/SELK-mediated NRF1 regulation offers profound clinical implications, particularly in overcoming drug resistance in cancer therapy. The NRF1-dependent cytoprotective program—which orchestrates de novo proteasome synthesis, aggrephagy (40)and ATF6-mediated upregulation of ERAD (41)—confers robust drug resistance to proteasome inhibitors. Furthermore, cancer cells exploit highly complex compensatory networks to maintain proteostasis, such as the stabilization of nuclear NRF1 via *O*-GlcNAcylation by OGT (42,43), and the complementary maintenance of basal proteasome activity and translational repression of NRF1 by the NRF3/CPEB3 axis (44). Because cancer cells employ these intricate layers of backup mechanisms, targeting the upstream ER-resident SELS/SELK machinery provides a highly synergistic strategy to completely disrupt this comprehensive “bounce-back” response and sensitize tumors to chemotherapy.

Conversely, in the context of neurodegenerative diseases characterized by toxic protein aggregation, modulating these ER-resident selenoproteins to safely promote NRF1 nuclear translocation could represent a promising therapeutic avenue. Such an approach would coordinately enhance proteasomal degradation, stimulate aggrephagy-dependent clearance of toxic aggregates, and alleviate ER proteotoxic stress (40,45). Future studies identifying the precise signaling cues—such as specific post-translational modifications or oxidative stressors—that switch the SELS/SELK machinery from maintaining NRF1 stability to facilitating its active retrotranslocation will further unmask the therapeutic potential of this intricate ER proteostasis network.

## 5. Conclusion

In conclusion, our study demonstrates that the ER-resident selenoproteins SELS and SELK cooperatively regulate the stability and stress-induced activation of the transcription factor NRF1. Rather than simply mediating the clearance of misfolded proteins, our results show that SELS and SELK act as functional regulators of NRF1 at the ER membrane. Specifically, SELS promotes the retention of NRF1 in the ER, whereas SELK induces its insolubilization. Furthermore, we found that the simultaneous loss of SELS and SELK leads to hyper-accumulation of nuclear NRF1 and excessive transcriptional activation of its target genes under proteasome impairment. These findings establish that SELS and SELK are essential for fine-tuning the basal stability and the appropriate “bounce-back” response of NRF1. Elucidating this SELS/SELK-mediated NRF1 regulation advances our fundamental understanding of ER quality control and provides a therapeutic rationale for targeting this machinery in human pathologies, such as overcoming proteasome inhibitor resistance in cancer or alleviating toxic protein aggregation in neurodegenerative diseases.

## Acknowledgements

We acknowledge the technical support of the Analytical Research Center for Experimental Sciences, Saga University. We also acknowledge Shinya Komoto of Optics and Imaging Facility, Okinawa Institute of Science and Technology Graduate University, and Motosuke Tsutsumi of National Institute for Physiological Sciences, National Institutes of Natural Sciences, for their technical assistance with the imaging.

## Authors’ contributions

G.Y., and T.T.: Conceptualization; G.Y., Y.K., T.O., R.Y., T.S., M.K., and T.T.: Methodology; G.Y., N.T., M.K., H.T., Y.S., Y.K., R.Y., T.S., and T.T.: Investigation; G.Y., and T.T.: resources; G.Y., N.T., M.K., H.T., Y.S., Y.K., R.Y., T.S., and T.T.: Data curation; G.Y., and T.T.: Writing original draft preparation; G.Y. and T.T.: Supervision; T.T.: Project administration; T.T.: Funding acquisition. All authors have read and agreed to the published version of the manuscript. This research was part of the dissertation submitted by the first author (G.Y.) in partial fulfilment of a Ph.D. degree. All authors have provided consent.

