## Supplemental Figure 1 for "Cooperative regulation of NF-E2 related factor 1 protein stability and transcriptional activation by endoplasmic reticulum-associated degradation system mediator, Selenoprotein S/K"

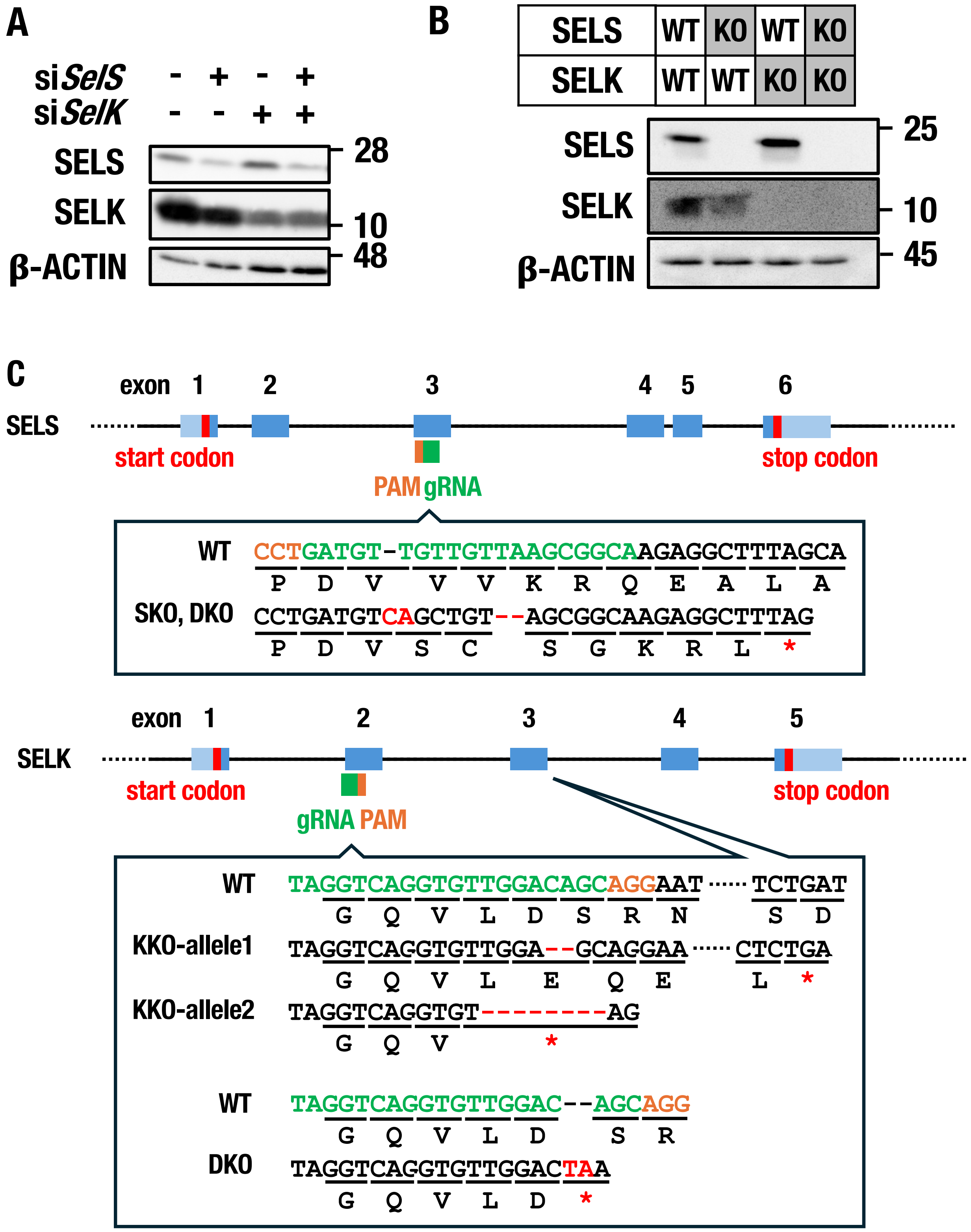


Supplemental Figure 1. Yamada et. al

**Supplemental Figure 1. Validation of SELS and SELK depletion in knockdown and knockout cell models.**

**(A) Confirmation of SELS and SELK knockdown.**

HEK293 cells are transfected with non-targeting control siRNA (siNT), *SelS* siRNA (si*SelS*), *SelK* siRNA (si*SelK*), or both *SelS* and *SelK* siRNAs (si*SelS/K*). Cell lysates were analyzed by immunoblotting to confirm the knockdown efficiencies of SELS and SELK.

**(B) Confirmation of SELS and SELK knockout.**

Cell lysates from wild-type (WT), SELS knockout (SKO), SELK knockout (KKO), and SELS/SELK double knockout (DKO) mouse embryonic fibroblasts (MEFs) were analyzed by immunoblotting to confirm the absence of SELS and SELK proteins.

**(C) Schematic representation of CRISPR-Cas9-mediated gene editing.**

The diagram illustrates the genomic locations of the guide RNAs (gRNAs) used for the generation of SKO, KKO, and DKO MEFs. Mutations introduced by the CRISPR-Cas9 system, including premature stop codons indicated by asterisks (*), nucleotide deletions indicated by dashes (–), and inserted bases, are all highlighted in red.
